# Do seabirds dream of artificial lights? Understanding light preferences of Procellariiformes

**DOI:** 10.1101/2024.03.01.582998

**Authors:** Elizabeth Atchoi, Mindaugas Mitkus, Biana Machado, Valter Medeiros, Sofia Garcia, Manuela Juliano, Joël Bried, Airam Rodríguez

**Affiliations:** Institute of Marine Sciences – OKEANOS, University of the Azores, Rua Professor Doutor Frederico Machado 4, 9901-862 Horta, Portugal; Institute of Biosciences, Life Sciences Center, Vilnius University, Vilnius, Lithuania; Ornithoscope Rogue Science – Faial, Portugal; Parque Natural de Ilha do Faial, Serviço de Ambiente da Ilha do Faial, Monte da Guia S/N, PT-9901-124 Horta, Açores, Portugal; Direção de Serviços de Biodiversidade e Política do Mar, Direção Regional de Políticas Marítimas, Secretaria Regional do Mar e das Pescas, Governo Regional dos Açores, Açores, Portugal, Rua D. Pedro IV, 29, PT-9900-111 Horta, Açores, Portugal; Marine Environment and Technology Laboratory-LAMTec-ID, University of Azores, Av. Álvaro Martins Homem Marina, Praia da Vitória Santa Cruz, 9760-412 Azores, Portugal; 8 Avenue de la reine Nathalie, 64200 Biarritz, France; Departamento de Ecología Evolutiva, Museo Nacional de Ciencias Naturales (MNCN), CSIC, Madrid, Spain

**Keywords:** Light pollution, Fallout, Ethology, ALAN, *Calonectris borealis*, color spectra

## Abstract

Seabirds, and particularly fledglings of burrow-nesting species, are greatly impacted by light pollution. During their inaugural flights from colony to sea, fledglings become grounded after encountering artificial light. Such groundings, or fallout events, affect many fledglings each year. To mitigate this light induced mortality, rescue programs have been implemented for decades in many locations worldwide. Despite the notoriety of fallouts, the contributing behavioural and biological factors remain mostly unknown. How do the mechanisms of light attraction and light avoidance interact or how do they manifest in different groups (e.g., age, personality, populations), or light pollution levels, remain open questions. We tested behavioural preferences of Cory’s shearwater *Calonectris borealis* fledglings, rescued after being grounded in urban areas, and of breeding adults, for contrasting light sources. Fledglings and adults were exposed to one of the three treatments in an experimental y-maze set-up: white light versus no-light, blue versus red light, and a control with no-light on each arm of the y-maze. Both age groups have shown preferences for the no-light arms and the red light arm. The preference for longer wavelengths and darker environments, along with slower responses by fledglings, suggest that close range light pollution appears to cause disorientation in seabirds. Our study helps to clarify the behavioural components of fallouts and provides further evidence on the disruptive effects of nocturnal artificial light on sensitive species.

## 1 Introduction

The increase of urbanization and development of new technologies continue to intensify anthropogenic pressures on ecosystems. The levels of artificial light at night at the Earth’s surface, or light pollution, are increasing at estimated rates from 2% to 9.6% a year (Kyba *et al*., 2017, 2023), seeping into protected and remote habitats (Garrett, Donald and Gaston, 2019) and continuously eroding nightscapes in marine and terrestrial realms (Gaston and Sánchez de Miguel, 2022; Marangoni *et al*., 2022). The pervasive impact is worsened by the recent widespread transition to LED technology, which is leading to increased emissions of short-wavelength (blue-rich) light (Sánchez de Miguel *et al*., 2021, 2022), which produces larger effects on wildlife (Longcore *et al*., 2018; Longcore, 2023).

Seabirds are among the most threatened avian groups (Dias *et al*., 2019) and burrow-nesting species (mainly Procellariiformes) are particularly vulnerable to light pollution (Rodríguez et al., 2019). At the end of the breeding season, thousands of burrow-nesting seabird fledglings fall to the ground on urban areas after encountering light-polluted areas during their initial flights from nest to sea (Rodríguez et al., 2017). Such fallout events have been documented and mitigated for decades through rescue programs, e.g. in Hawaii (Raine *et al*., 2017), Canaries (Rodríguez et al., 2023), Réunion (Chevillon *et al*., 2022) and Azores (Fontaine, Gimenez and Bried, 2011). In these programs’ citizens and specialists search for grounded birds, collect them and after inspection for health by trained staff the birds are released at sea intending on mitigating direct mortality. Nights with new moon, foggy conditions and strong inland winds increase the magnitude of fallouts (Rodríguez et al., 2017; Syposz et al., 2018; Telfer et al., 1987). Closeness of the source colony to the artificially lit area can lead to higher amount of grounded birds (Crymble et al., 2020; Rodríguez et al., 2015; Wilhelm et al., 2021), although fledglings can also be redirected to light polluted areas from substantial distances (Friswold *et al*., 2023). The amount of light pollution viewed by the fledglings during their flights to the ocean, or after they have already reached the ocean, also seems to influence the number of affected birds (Troy *et al*., 2013). However, proximate behavioural and biological causes of fallouts remain mostly unresolved, and it remains uncertain how other mechanisms, such as dispersal from the colony, may contribute to the pool of grounded birds (Brown et al., 2023; Rodríguez et al., 2017).

Behavioural responses of seabirds to light sources are complex, with evidence of interplay between attraction and avoidance. Fledglings are observed near light sources doing erratic flights, apparently unable to leave the overall lit environment via flight. Fledglings display increased tortuosity in their flights the closer they are to lit areas and the more intense the light pollution levels are (Rodríguez et al., 2022). Apart from fledglings’ fallouts, there are many documented events detailing adult seabirds being drawn to artificial lights or even crashing into them, for example in the collision and entrapment records from lighthouses, ships and other lit structures at sea (Montevecchi, 2006; Gjerdrum *et al*., 2021; Ryan, Ryan and Glass, 2021; Coleman *et al*., 2022). Similarly, adult little penguins *Eudyptula minor*, a burrow-nesting sphenisciforme, showed preference for lit paths on land over darker ones and for paths lit with shorter wavelengths (blue light) over longer wavelengths of light (red light), upon returning to their colony (Rodríguez et al., 2018). On the other hand, adult Manx shearwaters *Puffinus puffinus*, when in flight above the colony, temporarily avoided the area whenever a light was turned on, especially during exposure to blue-rich light (Syposz *et al*., 2021). Likewise the sporadic presence of light from ships facing a colony of Yelkouan shearwater *Puffinus yelkouan* was associated with fewer adults coming to the nests during the lit periods (Austad *et al*., 2023). The disparity of behaviours recorded suggests that reactions to artificial light may vary because of intrinsic qualities of the bird, such as species, age, body condition, personality, experience or phenology, or the characteristics of artificial light, such as distance to the light source, its intensity, emission spectra, temporal and spatial extent, and contrast to the surrounding. So, to effectively manage the effects of light pollution on seabirds, it is essential to understand their behaviours towards light at different life stages and under different scenarios.

In this study, we used a two-choice experimental y-maze set-up to investigate behavioural preferences to light stimuli at two life stages (fledglings and adults) of a burrow-nesting seabird, Cory’s shearwater (*Calonectris borealis*). We aimed to clarify behaviours, within a close range proximity to artificial light sources, simulating grounding: (1) first we tested preferences of fledglings towards different light sources (white versus no light, and red versus blue), and (2) second, we tested if adults behaviour could explain the age bias observed in fallouts.

It is generally considered that blue-rich light is more disruptive to organisms (Falchi *et al*., 2011; Longcore *et al*., 2018), having led to a higher magnitude of behavioural responses. In experiments with different taxa (sea turtles, Witherington & Bjorndal, 1991; migratory passerines, Doppler et al., 2015; Zhao et al., 2020; and seabirds, Rodríguez et al., 2018; Syposz et al., 2021). Thus, we expected both fledglings and adults to mirror these tendencies and show a preference for corridors with no light or lit with longer wavelength (i.e., less disruptive light). Fallouts affect age groups disproportionately; fledglings are grounded by the thousands, during their first flights from nest to sea, whereas adults, which fly over lit areas when travelling to and from their colonies throughout the breeding season, are seldom found grounded (Rodríguez et al., 2017a), and can breed successfully in inland colonies (Chevillon *et al*., 2022). Thus, because adults have previous experience with light-polluted landscapes, we expected them to react faster to the experimental stimuli.

## 2 Methods

### 2.1 Study species

Cory’s shearwaters breed in subtropical north Atlantic archipelagos, with their largest population located in the Azores archipelago (BirdLife International, 2023). The breeding period spans from mid-February to early November, during which breeding adults nest in burrows and raise a single chick (Monteiro, Ramos and Furness, 1996). Fledging occurs in late October early November, when an estimated 6% of fledglings are annually grounded in fallout events in the Azores (Fontaine, Gimenez and Bried, 2011; Rodrigues *et al*., 2012). To minimise mortality, SOS Cagarro rescue campaigns have been conducted in the Azores each year since 1995. During these campaigns, grounded fledglings are collected and released at sea, reducing direct mortality caused by light pollution (Fontaine, Gimenez and Bried, 2011; Atchoi *et al*., 2021).

In our study, we sampled 131 fledglings and 84 adults. Fledglings were sampled from the pool of rescued individuals during the SOS Cagarro campaign on Faial Island, Azores. All rescued individuals were brought to the operational centre in Horta, the main town of the island. We selected a cut-off minimum weight of the fledglings to ensure each individual was sufficiently healthy to take part in the experiment and be released thereafter. The minimum for selection was 660 g, and the mean for selected studied fledglings was 820 ± 74 g (± standard deviation, n = 128). The range of weight of fledglings rescued during 2020 and 2021 SOS campaign was 360 to 1080 g, average 766 ± 96 g (n = 1111), comparable to the long term range average of 750g (Cuesta-García *et al*., 2022). We selected 660 g to account for the number of available birds but still ensuring healthy weights to survive release. We ringed each with a unique numbered metal band and took biometrics of the fledglings in the mornings following capture, then placed the birds into rescue boxes (cardboard boxes of 20 × 20 × 40 cm with ventilation holes used to temporarily keep rescued birds during the SOS Cagarro campaign) and left in a quiet and dark place until the experimental test, which occurred in the evening of the same day. We tested fledglings for five nights in 2020 and for four nights in 2021, during October and November (Table S1). The experimental maze was placed on the edge of a 20-m high cliff (38.5199N, 28.6202W) on the outskirts of Horta, Faial Island, Azores. The two exits of the maze, which were parallel to each other, were set perpendicular to the cliff face, oriented at 100º E, facing the sea. Fledglings exiting the maze would have a small drop (< 50 cm) to a grassy slope, from where they could fly towards the sea whenever ready.

Adult shearwaters were captured and tested at the breeding colony of Capelinhos (38.5895N, –28.8233W), a Special Protection Area (SPA) from the Natura 2000 Network, an area with low light pollution levels situated on Faial Island. Capture occurred after sunset, either at the nest or on the ground. Each adult was ringed with a unique numbered metal band and had its biometric measurements taken. After capture and ringing, adults were transferred into SOS carboard boxes and brought to the testing site, just at the foot of the colony (< 20m), where they were kept for a minimum of 15 minutes before being tested to give the birds a time period to adjust to the darkness and adjust after the human manipulation. We tested adults for eleven nights between May and September 2022 (Table S1). The maze was placed the foot of the colony, in a flat ground-levelled area facing the sea, within < 10 m from the ocean surf. Adults leaving the maze walked into the colony grounds and were gently redirected to abandon the area directly in front of the maze to avoid choice bias for the next bird tested.

Handling and data collection from Cory’s shearwaters and the execution of the experiment were conducted under the licences from the Azorean government 46/2020/DRA, 60/2021/DRAAC, 48/2022/DRAAC, 25/2021/DRCTD and 36/2022/DRCTD, and the ethical licenses from University of the Azores UAC/2020/4182, UAC/2021/5720. Biometric data from grounded Cory’s shearwater fledglings (2020-2021) were collected by EA in the frame of the SOS Cagarro Campaign, coordinated by the Regional Directorate for Maritime Policies (DRPM of the Azorean Regional Government), with the collaboration of a network of institutional partners, including the Faial Island Environment and Climate Change Service. The responsibility for all conclusions drawn from the data lies entirely with the authors.

### 2.2 Experimental procedure

#### 2.2.1 Maze structure

The two-choice maze was a 100 x 50 x 40 cm plywood box, painted in matte black, with a separator at the middle to create two distinct exits (Figure S1). Lights were placed on the top of the maze in such a way that allowed each to illuminate the corresponding chamber as well as the bird from above. A smaller wooden box (black box) with black painted inner walls was fitted inside the back of the maze (Figure S1). This box was able to move vertically, so that we were able to cover the bird inside it and pull the box upward to release the bird into the maze revealing the choice arms. At its highest position, when the bird would be free to move around the maze, the walls of the box limited light spill from one side to the other. Individuals were placed one at a time inside the black box, at a centre position in the back area of the maze and stayed under the cover of the black box for one minute. After one minute the black box was pulled upward where it stayed for the duration of the test. Behaviours were directly observed via the observation window at the back of the maze and by visual confirmation at the choice arms whenever an individual exited the maze. To avoid confounding effects of odour, the maze was regularly cleaned with 70% ethanol (Bonadonna *et al*., 2006) after a couple of tests, or whenever an individual had defecated or regurgitated inside the maze. A black cardboard sheet was placed at the back of the maze where the individuals stood at the initial placement and replaced whenever the individual defecated or at the end of each experimental night.

#### 2.2.2 Light stimuli

For the white light stimulus, we used a commercially available LED strip (3000K warm white; AA-20-FLEX; by LEDsupply) with a broad emission spectrum with one blue peak at 455 nm and a second and highest at 625.2 nm (Figure S3). This light was not adjusted to be isoluminant for the shearwaters’ visual system as were the red and blue stimuli, because it was not to be tested in pair with either red or blue LEDs. The white light type was chosen to be a close simulation of the urban areas these fledglings and adults encounter, i.e., new installations of LED lights in Azorean urban areas which range from 3000K to 4000K (Electricidade dos Açores personal communication). After being installed in the maze, the white LEDs yielded an illuminance of 175 lux at the starting position of a bird. During control trials the illuminance at the birds starting position was > 0.1 lux which is the minimum detection range of the light meter used. We detected ∼175 lux in the white area floor and > 0.1 lux in the dark arm; corresponding to 2 cd/m^2^ reading a value of 20 cd/m^2^over white paper, and >0.1 on the dark area, both on the ymaze floor as in the white paper reading

The red and blue LED light systems used in this study were the same as in Atchoi et al., 2023. Briefly, the spectral sensitivities were modelled based on visual parameters of the wedge-tailed shearwater *Ardenna pacifica*, the most closely related species to Cory’s shearwater for which such data exist. The red and blue lights were designed to be as isoluminant as possible for the achromatic visual system, but to have strong contrast for the chromatic visual system. The final adjustment of the light stimuli yielded a strong chromatic contrast of 60.7 JNDs (just noticeable differences; the discrimination threshold is 1 JND) between them. The achromatic contrast between the two stimuli was 4%. As most birds tested to date have very poor achromatic contrast sensitivity to stationary stimuli (usually birds cannot discriminate contrast below 10% or even more (Lind *et al*., 2012; Olsson, Lind and Kelber, 2018)) we believe that Cory’s shearwaters would not be able to discriminate our stimuli based on luminance alone. For human visual system, the blue LEDs yielded an illuminance of 75 lux and the red LED of 120 lux at the starting position of a bird (measured with a Hagner Screen-Master light meter; B. Hagner, Solna, Sweden). These illuminances correspond to light levels at civil twilight (Cronin T. W., 2014) and are comparable to what a chick would be exposed to outside its burrow after sunset. Further details on the red and blue light system, including dimensions, measurements, and sensitivity modelling, can be found in Atchoi et al., 2023, and in the supplemental material (Figure S2).

#### 2.2.1 Treatments and variables

We used a two-choice maze to test the behavioural preferences of fledglings and adults against light stimuli. Each bird was subjected to only one trial, and presented with one out of three treatments: (1) White-Dark, where one arm had no light stimulus (dark), and the other was fitted with a white light; (2) Blue-Red, where one arm was illuminated by a red light and the other by a blue light of equal intensities (see “Light stimuli” above); and (3) Control, where both arms were without any experimental light, i.e., both choices were ‘dark’, but to be able to identify any bias generated by the structure of the box itself, we registered if the bird chose ‘left’ or ‘right’ arm. To eliminate physical bias of the maze structure itself, the position of the lights were randomly interchanged between arms. For each bird, we recorded (1) choice (yes or no), (2) preference (white or dark, red or blue, left or right) and (3) the latency to make a choice (in minutes). A choice was registered if a bird stepped into one of the arms of the maze, regardless of whether it exited the maze or not. No-choice was registered if a bird stayed around the starting position, even if it was facing or near one of the arms (Figure S1). Whenever an individual made no choice, the test ended after 10 minutes and the individual was removed from the box and released. Because the treatments were assigned in a random order it resulted in an uneven number of trials per treatment (Figure 1).

**Figure 1.**
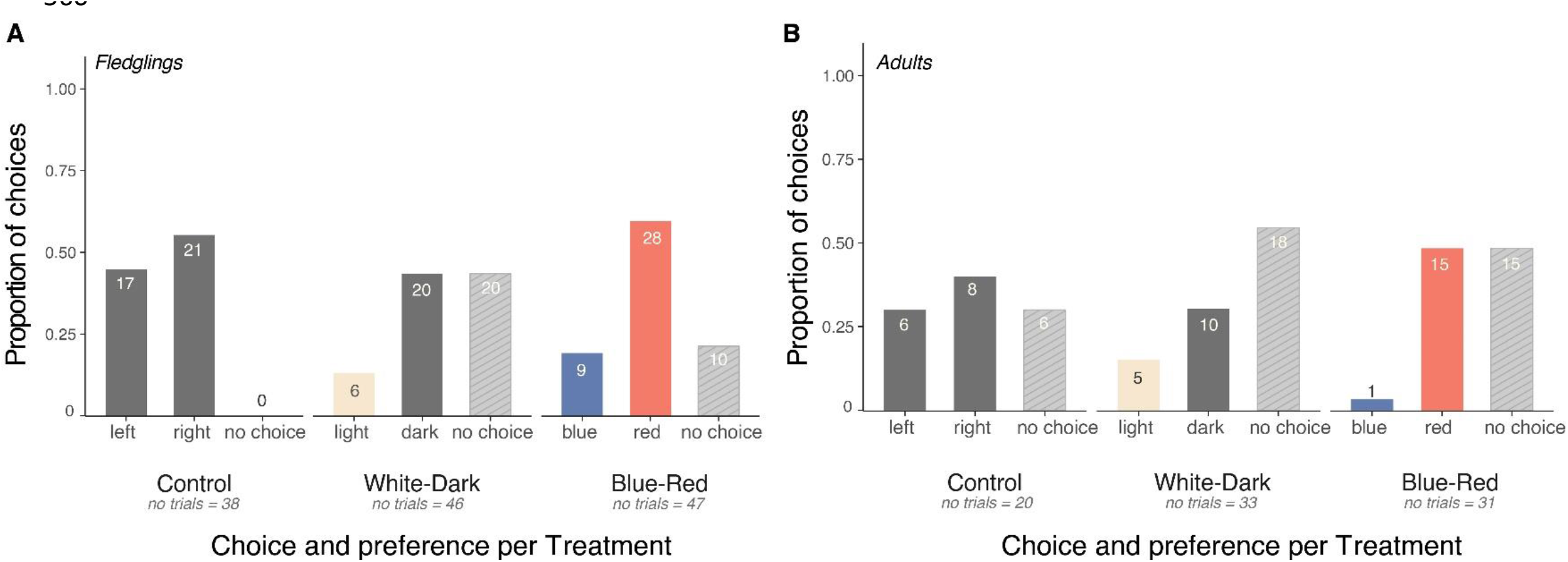
count of behaviours and preference per treatment for (A) 131 fledglings and (B) 84 adults. Striped columns represent no-choices. Number on bars indicate the number of individuals.

### 2.3 Statistical analysis

We ran a Fisher exact test to determine the differences between the two age groups concerning the number of choices and the type of choices made. Preference was analysed with an exact two-tailed binomial test, for each treatment separated by age group. To estimate differences in latency to make a choice within treatments for each age group we used Welch two sample t tests. Since a single measurement may not reflect body size in a reliable manner (Rising and Somers, 1989), we conducted a principal component analysis (PCA) on biometric data (wing, tarsus, culmen and gonys (mm); Figure S4). We then used the estimates from the first component (which retained 51% of the variation) as a body size index (BSI), following Rodríguez Dann & Chiaradia 2017.The relationship between PC1 and biometrics was negative, so we multiplied BSI by –1 for a better interpretation. Subsequently we calculated body condition indices (BCI) as the standardized residuals of a linear regression of body mass on BSI. Positive values of BSI and BCI respectively represent larger individuals, and individuals heavier than average after controlling for body size, i.e. in a presumably better body condition (heavier fledglings have higher survival rates; Mougin et al., 2000).

Generalized linear models with binomial error distributions and logit link functions were conducted for each age group and each treatment to detect relationships between continuous explanatory variables (BSI and BCI), and the binomial response variables a) choice (yes = 1, no = 0) and b) preference (white = 1, dark = 0; blue = 1, red = 0; left = 1, right = 0). Linear models were conducted to determine relationship between the continuous explanatory variables (BSI and BCI) and a continuous variable (time) for the latency to choice. All statistical analyses and data manipulation were conducted using R version 4.1.2. (R Core Team, 2021).

## 3 Results

### 3.1 Choice or no choice

Fledglings were overall 2.88 times more likely to make a choice, making choices in 77% of trials (101/131) while adults did so in 54% of trials (45/84) (Fisher exact test: p = < 0.001, Odds Ratio = 2.88; Table S2) with differences being detected among treatments (Figure 1). In the White-Dark treatment both groups showed similar choice rates (Fisher exact test: p = 0.367, Odd Ratio = 1.55)(Table S2). Fledglings made more choices than adults in the Blue-Red treatment (Fisher exact: p = 0.015, Odds Ratio = 3.4). During Control treatment all fledglings made a choice, while adults only made a choice in 70% of the trials.

### 3.2 Preference

During the White-Dark treatment fledglings consistently chose the dark arm of the maze over the lit one, whereas adults showed a less definitive preference, despite also choosing the dark arm more often (Figure 1; binomial exact test, fledglings p = 0.009; adults p = 0.3; Figure 2). Both age groups showed a clear preference for red light during the Blue-Red treatment (Figure 1; binomial exact test, fledglings p = 0.003; adults p = 0.00052; Figure 2). In the Control treatment there was no difference between left and right arm choices for either age group (binomial exact tests, fledglings: p = 0.627; adults: p = 0.115; Figure 2).When comparing preferences between age groups, we found that fledglings were as likely as adults to choose the red and dark arms of the maze (Fisher exact test, red light: p = 0.25; dark: p = 0.5).

**Figure 2.**
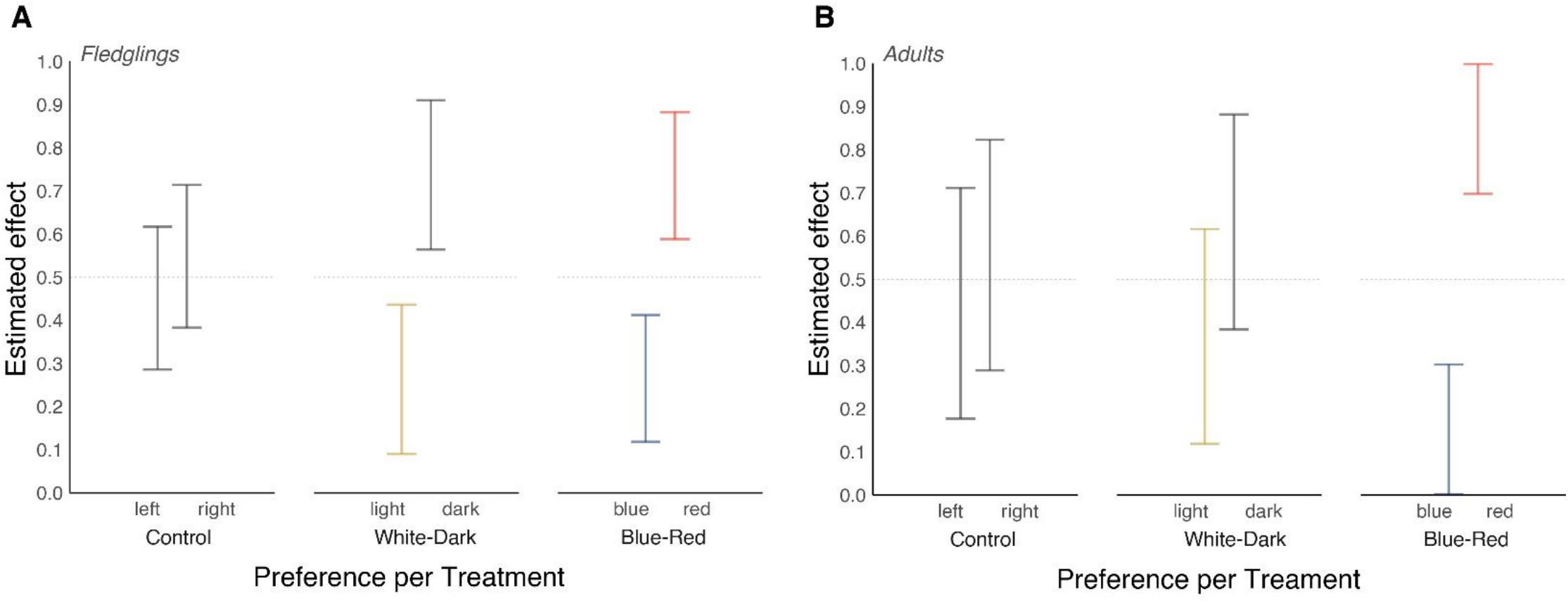
Results of exact binomial two-tailed tests for each preference per treatment. (A) data for fledglings and (B) data for adults. Bars represent 95% confidence intervals. Significant differences where 95% CIs do not intercept 0.5.

### 3.3 Latency to make a choice

Fledglings were slower to make a choice than adults (Figure 3). Both age groups were quicker in Control treatment than in any of the other two treatments (Figure 3). Fledglings’ latency to make a choice did not differ regardless of choosing white-light or dark arm during the White-Dark treatment. Fledglings appeared to be quicker when choosing the lit arm yet the differences were not statistically significant (Welch two sample t test: t_7_ = –0.989, p = 0.353; Figure 3). Latency to choice also did not differ between blue or red arm in the Blue-Red treatment (Welch two sample t-test: t_15_ = –1.04, p = 0.314) or whether they chose the left or the right arm of the maze in the Control treatment (Welch two sample t test: t_23_ = 0.65, p = 0.521).

**Figure 3.**
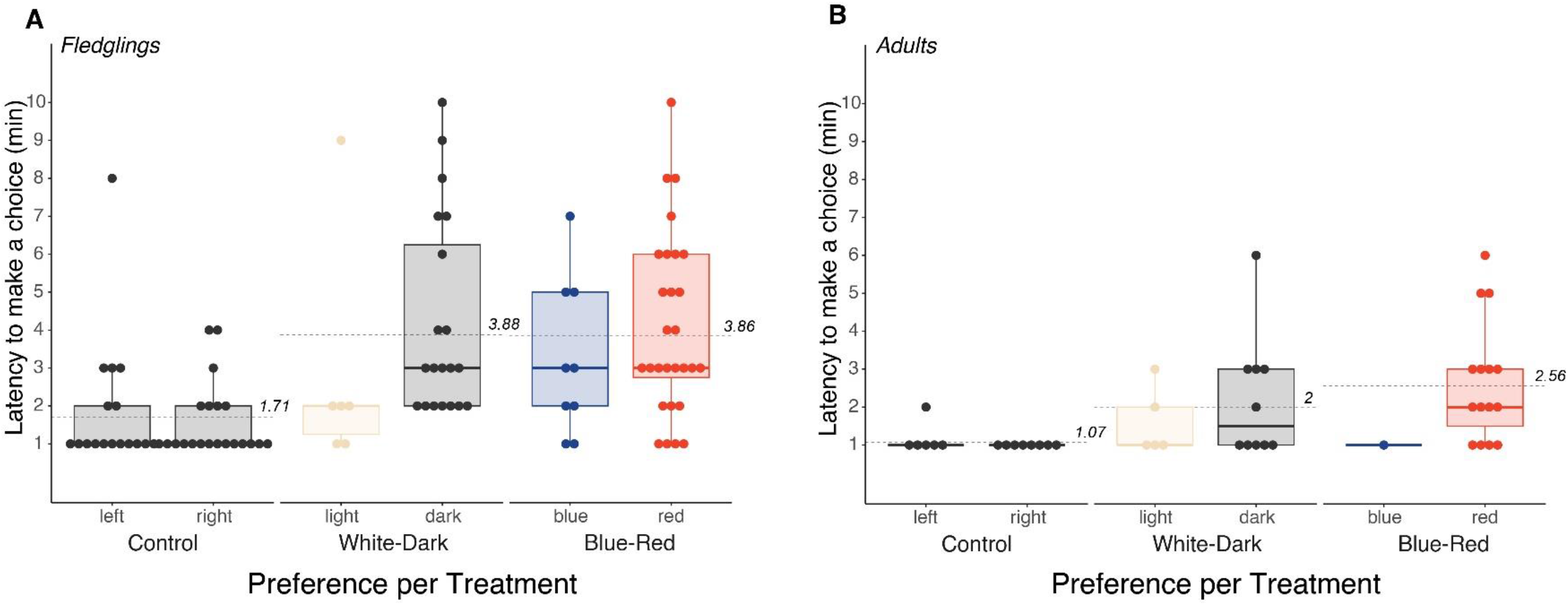
Latency to choose by Treatment and choice. A) fledglings and B) adults. Means for each treatment in dashed line with corresponding value.

Adults’ latency to make a choice did not differ between lit or dark arm during White-Dark treatment (Welch two sample t test: t_12_ = –0.923, p = 0.373), nor did it differ between left or right arm during the Control treatment (Welsh two sample t test: t_5_ = 1, p = 0.363). It was not possible to determine differences for adults’ latency of choice between blue and red as only one adult chose the blue arm

### 3.4 Body Size Index and Body Condition

Only two weak relationships were given by the models, first, larger fledglings showed a tendency to choose more often during Blue-Red treatment (glm estimate: 0.581, 95%CI = [0.007, 1.253]), and second, larger adults showed a tendency to choose more often during Control (glm estimate: 1.161, 95%CI = [0.120, 3.000]). No other significant relationship between BSI or BCI and preference or latency to choose were observed (Table S3).

## 4 Discussion

Our study verified behavioural preferences of Cory’s shearwaters for light stimuli and showed that these preferences do not differ significantly between fledglings and adults. However, age-specific differences in the frequency and time taken to make a choice were identified. Indeed, former studies showed that Cory’s shearwater chicks displayed a preference for dark environments when exposed to artificial light at a closer range (Atchoi *et al*., 2023), and adults of the closely related species, such as Manx and Yelkouan shearwaters, avoided their colony when it was sporadically exposed to artificial light (Syposz *et al*., 2021; Austad *et al*., 2023). Both age groups displayed stronger avoidance of shorter wavelength artificial lights.

### Effect of artificial light stimuli

In our study, all fledglings made a choice whenever light stimuli were absent from the experimental maze, i.e. in Control treatment. However, when light stimuli were present (White-Dark and Blue-Red treatments), some fledglings were deterred from making a choice (Figure 1). Furthermore, the fledglings that made a choice, were quicker to do so in Control trials, but slower in treatments with light stimuli (Figure 3). Our data indicate that presence of a close-range artificial light can be disruptive to fledglings of Cory’s shearwaters, for example, because of disorientation or fright, influencing their behaviour. More specifically, artificial light appears to impair the fledglings’ ability to exit the unknown lit environment, thus increasing their vulnerability to dangers present in urban areas for those who are grounded. Our results agree with the observations of irregular and chaotic flights displayed by fledglings during fallout events (pers. observation EA,TP,AR, BM, & JB). Attraction to light may be the proximate cause leading fledglings to converge on urban lit areas. However, when fledglings come into close range of artificial lights, they become apparently disoriented by the stimuli rather than being behaviourally attracted to them (see tracked flights in Rodríguez et al., 2022).

### The role of the visual system

Disorientation could be a result of artificial light interfering with the bird’s visual system. Studies have described an underdeveloped and untrained visual system in growing Procellariiformes chicks: older Leach’s-storm petrel chicks *Hydrobates leucorhous* reacted more times to light stimuli than younger ones, i.e., underdeveloped (Mitkus, Nevitt and Kelber, 2018), and growing Cory’s shearwater chicks reacted more to light stimuli the more they had been previously exposed to light, i.e., trained (Atchoi *et al*., 2023). In the current study, fledglings were slower to react than adults in general (Figure 3), suggesting that the light-induced effect is stronger for the younger birds. Additionally, the slower reactions of fledglings, even in Control treatments, indicates their general behavioural inexperience. Our results suggest that the latent visual system development in fledglings of burrow nesting species (Atchoi, Mitkus and Rodríguez, 2020) will increase the vulnerability to the adverse effects caused by light pollution upon fledgling, and that this vulnerability, observed by the apparent disorientation displayed by the flights of fledglings, is reduced in adults birds with both visual and navigational experience (Imber, 1975).

The delayed development of the visual system of underground nesting Procellariiformes could, in turn, be explained by the growing conditions of the chicks; chicks of these species spend the first weeks of their life in dark burrows which lack sufficient light stimuli needed for a proper development of a functioning visual system (Mitkus, Nevitt and Kelber, 2018; Atchoi *et al*., 2023). The interference of light with the visual ability of fledglings as they begin navigate in flight these new lit environment, might be a factor explaining the age bias observed in fallout event. Our data showed similar behavioural preference between age groups, so behavioural differences cannot account for the age bias observed in fallouts. Adults displayed similar behaviours to fledglings with overall fewer choices made during treatments with light stimuli (Figure 1) and being quicker to exit the maze during Control treatment. As with fledglings, the presence of artificial light also contributed to a disruption of the adults’ behaviour. Other studies observed adult procellariiforms avoiding their colonies when a temporary light source was turned on (shearwaters, Austad et al., 2023; Syposz et al., 2021).

### Effect of artificial light spectra

Both fledglings and adults presented similar preferences for the dark arm of the maze versus the white light and for the red versus the blue LED arm (Figure 1; Figure 2). The behavioural preferences displayed by adults in the White-Dark treatment were less distinct than the preferences of fledglings (Figure 1; Figure 2), even if the proportions were similar (fledglings: 23% chose white, 77% chose dark; adults: 33% chose white, 66% chose dark). The less clear preference by adults could be attributed to a smaller sample size (Figure 1), and the tendency should be confirmed if more adults were sampled. We executed a power analysis which indicated the need of between 176 and 305 birds to confirm the tendencies. Yet, the similarly low sample size used in the Blue-Red treatment was sufficient to result in clear preferences for both age groups (Figure 1; Figure 2).

Blue-rich light has been found to be more disruptive to burrow-nesting seabirds (Rodríguez et al., 2017a, 2018; Syposz et al., 2021), and most recommendations to reduce the overall effect of, and the amount of, light pollution highlight the shifting of urban light spectral emissions towards longer wavelengths (Gaston & Sánchez de Miguel, 2022; Rodríguez, 2023). Accordingly, in our study, we interpreted the preference for the red-light and dark arms of the maze as evidence for stronger effects induced by the blue-light, which is also presents in a broad spectrum white-light (it has a peak emission in the short wavelengths) (Figure S2; Figure S3). Indeed, procellariiforms forage over, and dive into, waters typically rich in short wavelengths (400-500 nm), accordingly their visual systems seem to be attuned to such environments (Hart, 2004), making individuals more sensitive to blue-rich artificial lights.

Notably, the White-Dark treatment had a more pronounced effect on adults (Figure *1*,Figure *2*). The white-dark treatment also revealed an increased reaction speed when fledglings chose white or otherwise a higher latency when choosing dark. Although we did not directly measure and compare emissions between the white and the blue or red LEDs for the shearwater colour visual model (see methods) (Figures S2 and S3), photometric measurements within the maze (in lux) suggests that the intensity of white light should have been similar to that of blue and red LEDs as they all share similar intensity magnitude (white 175 lux; blue 75 lux; red 120 lux). Despite this similarity, during the White-Dark treatment, fewer fledglings and adults made a choice compared to trials with the Blue-Red treatment, and even fewer than during the Control treatment (Figure 1). Furthermore, fledglings selecting the white-light arm exhibited much faster reactions than those choosing the dark arm (Figure 3). Given the comparable intensity of the three light treatments, we do not attribute the stronger effect on time to reaction? of the White-Dark treatment to differences in light intensity. Rather, the White-Dark treatment presented a more contrasting environment, with white light against a dark area (∼175 lux in the white area and > 0.1 lux in the dark arm, all values for illuminance (lux) and lluminance (cd/m^2^) can be seen in Fig. S4), compared to the blue versus red light (∼45 lux difference between both arms). This high contrast of the White-Dark treatment may weaken the birds’ decision-making capabilities and worsen disorientation caused by light pollution after grounding. We can extrapolate from these results an infer that high contrast between well-defined lit and dark zones may also disorient birds in flight (Guilford *et al*., 2018). The broad emission spectra of the white-light treatment may also contribute to a stronger response. Indeed, Manx shearwater adults showed a greater aversion to flying into the colony when exposed to white light compared to blue light (Syposz *et al*., 2021). Additionally, broad white light emissions led to more grounded fledglings compared to more monochromatic lights (Rodríguez, Dann and Chiaradia, 2017).

### Innate behavioural preferences

The fledglings tested in our study had already been grounded by artificial lighting in urban areas. These individuals could have been biased to avoid light after having already associated lit areas with stress. If individuals with limited or no exposure to artificial light during growth were tested just before being exposed to urban light, they might display distinct behaviours. In other words, unexposed shearwater fledglings might innately display a preference for light sources versus dark environments. Indeed, there is evidence of behavioural shifts in other cavity-nesting bird taxa, for example, in common starlings *Sturnus vulgaris*, which display negative phototaxis as young chicks, but switch to positive phototaxis as they approach fledging (Minot, 1988). However, previous work revealed avoidance of light stimuli in Cory’s shearwater chicks aged from three weeks old to nearly fledging (Atchoi *et al*., 2023), suggesting ours results were not skewed due to the previous exposure to artificial light at night, nor do Cory’s shearwaters go through a behavioural shift similar to that of starlings. Testing light-unexposed fledglings should further clarify the behavioural preferences of this species, yet the practical constraints associated with fieldwork make it challenging to obtain an adequate sample size.

### Vulnerability differences at the individual level

In our study not all birds made a choice, particularly adults (only half of them completed the trials) (Figure 1). While it is difficult to determine why individuals make or do not make choices in such settings, it has been proposed that a choice may be derived from specific needs at the time of testing (Brooke, 1998) or personality differences (Bonadonna and Sanz-Aguilar, 2012). In our Control treatment, 70% of adults made a choice, whereas all fledglings made a choice. This agrees with the assumption that fledglings are driven by hunger and an instinct to reach the ocean and forage (fledglings lose ca. 40 g per day until the first successful foraging event (Mougin *et al*., 2000)). On the other hand, the behavioural preferences of healthy adults should be activity-dependent.

The link between personality and body condition is poorly understood, with inconclusive results (Réale *et al*., 2007). While body condition could affect behavioural responses to stress (Seltmann *et al*., 2012), our results did not yield consistent patterns regarding body size, body condition, and changes in behaviour (Figure 4; Figure 5). Our models detected only two weak effects (see estimates and confidence intervals in Table S3).

**Figure 4.**
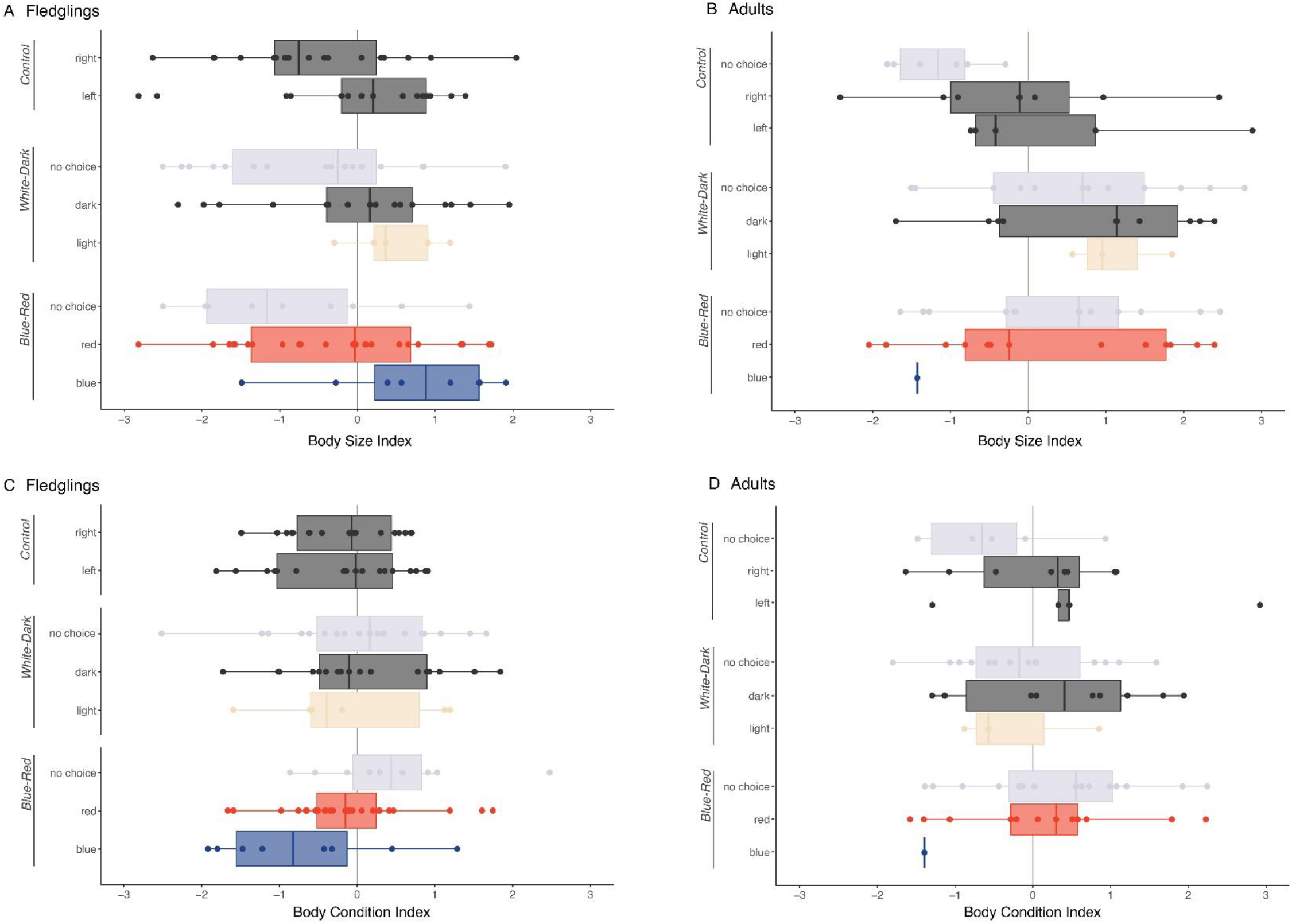
Body size of (A) fledglings and (B) adults, and Body condition index of (C) fledglings and (D) adults per behaviour and trial. All fledglings made a choice during Control trials, thus only two boxplots in (A) and (C) for control treatment.

**Figure 5.**
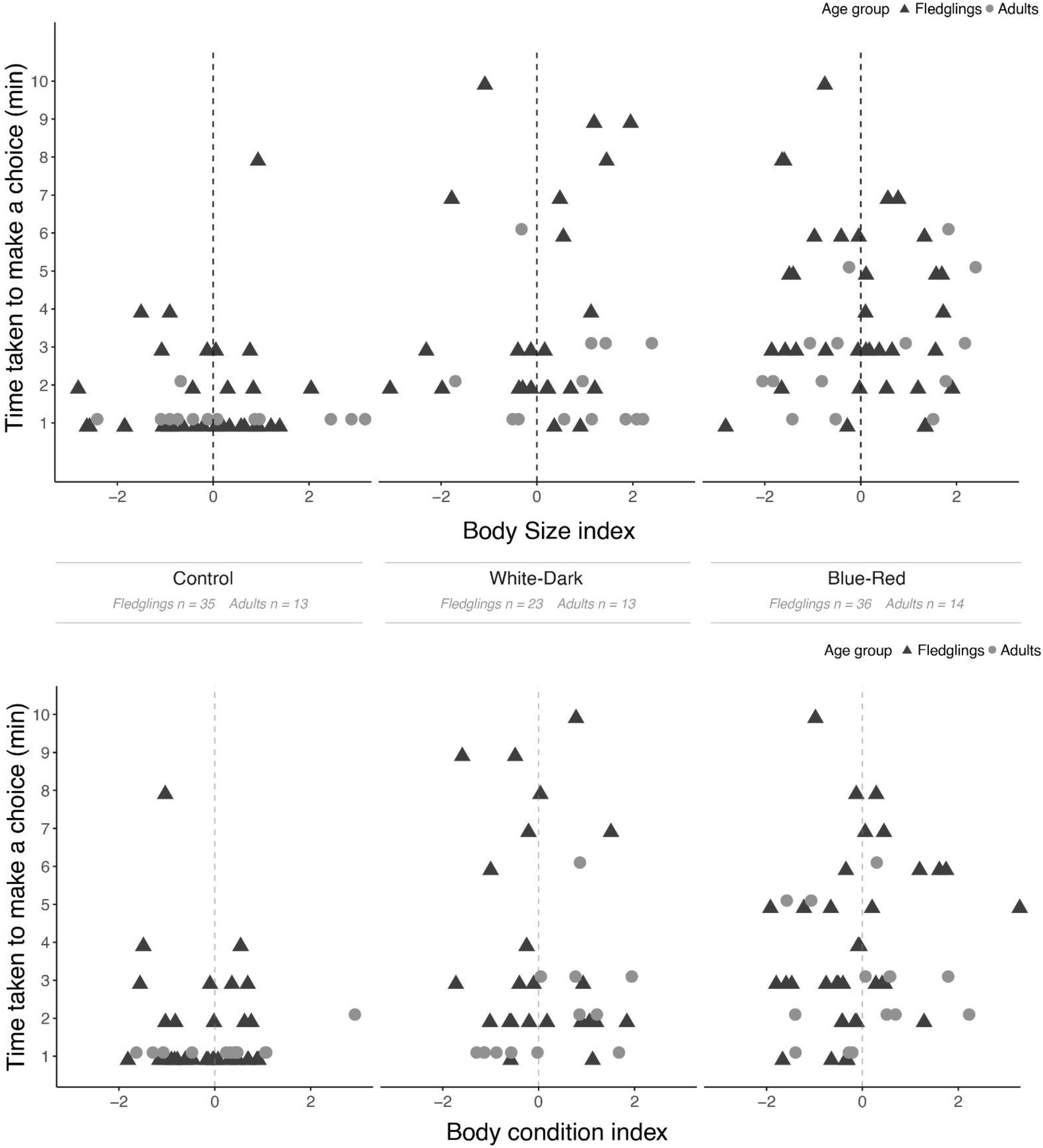
Latency to make a choice against body size and body condition per age group and trial type. Vertical dashed grey line represents an average body condition (value = 0).

We sampled fledglings based on their apparent health and heavier body mass (see Methods), limiting the representativeness of the sample. Random sampling of grounded fledgling without body mass thresholds, could unveil relationships between body condition and behaviour towards light stimuli undetected in this study. The consequences of exhibiting bolder or shyer behaviour in the context of fallout events should be further investigated.

Finally, other factors, such as differential exposure to light during growth, via the amount of natural or artificial light that enters a burrow, might lead to differences in visual system development (Atchoi *et al*., 2023) or even personality (Ruiz-Raya, Noguera and Velando, 2022). These factors might also partially explain the rate of responses and vulnerability to light pollution at the individual level. Further research is needed to identify the most vulnerable subgroups, and to determine how mitigative measures can be extended in both time and space to reduce the effects of light pollution on seabird’ populations.

## Acknowledgements

We are grateful to Joseph Lewin for constructing the maze and Lorenzo Kunze for providing expert logistical support to the light stimuli. We thank Pranciškus Vitta, Olle Lind and Peter Olsson for the help and advice in preparation of light stimuli. We thank OMA (Observatório do Mar dos Açores) for providing personnel and support during SOS Cagarro campaign and field work. We thank volunteers João Lagoa, Maria São Bento and Hannah Gretham for contributing to the data collection, and Ellie Ga for the constructive discussions on the topic which greatly improved the study. Thanks to volunteers Pedro Marques and Alberto Lopez for collecting data during SOS Cagarro campaign and improving the discussion of the topic, and to Scott Cameron at LEDsupply for technical guidance and general notes on seabird welfare. We are grateful to Faial Island Environment and Climate Change Services and its rangers for the logistical support. Finally, the authors extent their gratitude to the Regional Directorate for Maritime Policies (DRPM of the Azorean Regional Government) for providing access to the rescued fledglings and to all volunteers and trained staff involved in the collection, rescue and processing of grounded Cory’s shearwaters during the SOS Cagarro campaign on Faial Island. Rescued fledglings’ biometrics were collected as part of the SOS Cagarro Campaign, coordinated by the Regional Directorate for Maritime Policies with the collaboration of a network of institutional partners, including the Island Environment and Climate Change Services. The responsibility for all conclusions drawn from the data lies entirely with the authors.

## References

Atchoi, E. et al. (2021) ‘LuMinAves: cooperative research and mitigation of light pollution impacts in seabirds’, International Journal of Sustainable Lighting, 23(1), pp. 33–41. doi: 10.26607/ijsl.v23i1.107.

Atchoi, E. et al. (2023) ‘Ontogenetic exposure to light influences seabird vulnerability to light pollution’, Journal of Experimental Biology, p. jeb.245126. doi: 10.1242/jeb.245126.

Atchoi, E., Mitkus, M. and Rodríguez, A. (2020) ‘Is seabird light-induced mortality explained by the visual system development?’, Conservation Science and Practice, 2(6), p. e195. doi: 10.1111/csp2.195.

Austad, M. et al. (2023) ‘The effects of temporally distinct light pollution from ships on nocturnal colony attendance in a threatened seabird’, Journal of Ornithology, (0123456789). doi: 10.1007/s10336-023-02045-z.

Bonadonna, F. et al. (2006) ‘Evidence that blue petrel, Halobaena caerulea, fledglings can detect and orient to dimethyl sulfide’, Journal of Experimental Biology, 209(11), pp. 2165–2169. doi: 10.1242/jeb.02252.

Bonadonna, F. and Sanz-Aguilar, A. (2012) ‘Kin recognition and inbreeding avoidance in wild birds: The first evidence for individual kin-related odour recognition’, Animal Behaviour, 84(3), pp. 509–513. doi: 10.1016/j.anbehav.2012.06.014.

Brooke, M. D. L. (1998) ‘Determination of the absolute visual threshold of a nocturnal seabird, the Common Diving Petrel Pelecanoides urinatrix’, ibis, (131), pp. 290–300.

Brown, T. M. et al. (2023) ‘A path forward in the investigation of seabird strandings attributed to light attraction’, Conservation Science and Practice, 5(1), p. e12852. doi: 10.1111/csp2.12852.

Chevillon, L. et al. (2022) ‘25 years of light-induced petrel groundings in Reunion Island: Retrospective analysis and predicted trends’, Global Ecology and Conservation, 38(June). doi: 10.1016/j.gecco.2022.e02232.

Coleman, J. et al. (2022) ‘Blinded by the light: Seabird collision events in South Georgia’, Polar Biology, (0123456789). doi: 10.1007/s00300-022-03045-0.

Cronin T. W. (2014) Visual ecology. Princeton University Press.

Crymble, J. et al. (2020) ‘Identifying light-induced grounding hotspots for Maltese seabirds’, Il-Merill, 34, pp. 23–43.

Cuesta-García, M. et al. (2022) ‘Targeting efforts in rescue programmes mitigating light-induced seabird mortality: First the fat, then the skinny’, Journal for Nature Conservation, 65(October 2021). doi: 10.1016/j.jnc.2021.126080.

Dias, M. P. et al. (2019) ‘Threats to seabirds: A global assessment’, Biological Conservation, 237(June), pp. 525–537. doi: 10.1016/j.biocon.2019.06.033.

Doppler, M. S. et al. (2015) ‘Cowbird responses to aircraft with lights tuned to their eyes: Implications for bird-aircraft collisions’, Condor, 117(2), pp. 165–177. doi: 10.1650/CONDOR-14-157.1.

Falchi, F. et al. (2011) ‘Limiting the impact of light pollution on human health, environment and stellar visibility’, Journal of Environmental Management, 92(10), pp. 2714–2722. doi: 10.1016/j.jenvman.2011.06.029.

Fontaine, R., Gimenez, O. and Bried, J. (2011) ‘The impact of introduced predators, light-induced mortality of fledglings and poaching on the dynamics of the Cory’s shearwater (Calonectris diomedea) population from the Azores, northeastern subtropical Atlantic’, Biological Conservation, 144(7), pp. 1998–2011. doi: 10.1016/j.biocon.2011.04.022.

Friswold, B. et al. (2023) ‘From colony to fallout: Artificial lights pose risk to seabird fledglings far from their natal colonies’, Conservation Science and Practice, (June), pp. 1–14. doi: 10.1111/csp2.13000.

Garrett, J. K., Donald, P. F. and Gaston, K. J. (2019) ‘Skyglow extends into the world’s Key Biodiversity Areas’, Animal Conservation, 23(2), pp. 153–159. doi: 10.1111/acv.12480.

Gaston, K. J. and Sánchez de Miguel, A. (2022) ‘Environmental Impacts of Artificial Light at Night’, Annual Review of Environment and Resources, pp. 1–26.

Gjerdrum, C. et al. (2021) ‘Bird strandings and bright lights at coastal and offshore industrial sites in Atlantic Canada’, Avian Conservation and Ecology, 16(1).doi: 10.5751/ACE-01860-160122.

Guilford, T. et al. (2018) ‘Light pollution causes object collisions during local nocturnal manoeuvring flight by adult Manx Shearwaters Puffinus puffinus’, Seabird, 31, pp. 48–55.

Hart, N. S. (2004) ‘Microspectrophotometry of visual pigments and oil droplets in a marine bird, the wedge-tailed shearwater Puffinus pacificus: topographic variations in photoreceptor spectral characteristics’, Journal of Experimental Biology, 207(7), pp. 1229–1240. doi: 10.1242/jeb.00857.

Imber, M. J. (1975) ‘Behaviour of petrels in relation to the Moon and artificial lights’, Notornis, 22(Gould 1967), pp. 302–306.

Kyba, C. C. M. et al. (2017) ‘Artificially lit surface of Earth at night increasing in radiance and extent’, Science Advances, 3(11), pp. 1–9. doi: 10.1126/sciadv.1701528.

Kyba, C. C. M. et al. (2023) ‘Citizen scientists report global rapid reductions in the visibility of stars from 2011 to 2022’, Science, 379(6629), pp. 265–268. doi: 10.1126/science.abq7781.

Lind, O. et al. (2012) ‘Luminance-dependence of spatial vision in budgerigars (Melopsittacus undulatus) and Bourke’s parrots (Neopsephotus bourkii)’, Journal of Comparative Physiology A: Neuroethology, Sensory, Neural, and Behavioral Physiology, 198(1), pp. 69–77. doi: 10.1007/s00359-011-0689-7.

Longcore, T. et al. (2018) ‘Rapid assessment of lamp spectrum to quantify ecological effects of light at night’, Journal of Experimental Zoology Part A: Ecological and Integrative Physiology, 329(8–9), pp. 511–521. doi: 10.1002/jez.2184.

Longcore, T. (2023) ‘A compendium of photopigment peak sensitivities and visual spectral response curves of terrestrial wildlife to guide design of outdoor nighttime lighting’, Basic and Applied Ecology, 73(October), pp. 40–50. doi: 10.1016/j.baae.2023.09.002.

Marangoni, L. F. B. et al. (2022) ‘Impacts of artificial light at night in marine ecosystems—A review’, Global Change Biology, 28(18), pp. 5346–5367. doi: 10.1111/gcb.16264.

Mitkus, M., Nevitt, G. A. and Kelber, A. (2018) ‘Development of the visual system in a burrow-nesting seabird: Leach’s Storm Petrel’, Brain, Behavior and Evolution, 91(1), pp. 4–16. doi: 10.1159/000484080.

Monteiro, L. R., Ramos, J. A. and Furness, R. W. (1996) ‘Past and present status and conservation of the seabirds breeding in the Azores archipelago’, Biological Conservation, 78(3), pp. 319–328. doi: 10.1016/S0006-3207(96)00037-7.

Montevecchi, W. A. (2006) ‘Influences of artificial light on marine birds’, in Longcore, T. and Rich, C. (eds) Ecological Consequences of Artificial Night Lighting. Island Press, pp. 94–113.

Mougin, J. L. et al. (2000) ‘Fledging weight and juvenile survival of Cory’s Shearwaters Calonectris diomedea on Selvagem Grande’, Ringing and Migration, 20(2), pp. 107–110. doi: 10.1080/03078698.2000.9674231.

Olsson, P., Lind, O. and Kelber, A. (2018) ‘Chromatic and achromatic vision: Parameter choice and limitations for reliable model predictions’, Behavioral Ecology, 29(2), pp. 273–282. doi: 10.1093/beheco/arx133.

Raine, A. F. et al. (2017) ‘Declining population trends of Hawaiian Petrel and Newell’s Shearwater on the island of Kaua’i, Hawaii, USA’, Condor, 119(3), pp. 405–415. doi: 10.1650/CONDOR-16-223.1.

Réale, D. et al. (2007) ‘Integrating animal temperament within ecology and evolution’, Biological Reviews, 82(2), pp. 291–318. doi: 10.1111/j.1469-185X.2007.00010.x.

Rising, J. and Somers, K. (1989) ‘The Measurement of Overall Body Size in Birds’, The Auk, 106(4), pp. 666–674. doi: 10.1093/auk/106.4.666.

Rodrigues, P. et al. (2012) ‘Remote sensing to map influence of light pollution on Cory’s shearwater in São Miguel Island, Azores Archipelago’, European Journal of Wildlife Research, 58(1), pp. 147–155. doi: 10.1007/s10344-011-0555-5.

Rodríguez, A. et al. (2017) ‘Seabird mortality induced by land-based artificial lights’, Conservation Biology, 31(5), pp. 986–1001. doi: 10.1111/cobi.12900.

Rodríguez, A. et al. (2018) ‘Penguin colony attendance under artificial lights for ecotourism’, Journal of Experimental Zoology Part A: Ecological and Integrative Physiology, 329(8–9), pp. 457–464. doi: 10.1002/jez.2155.

Rodríguez, A. et al. (2019) ‘Future directions in conservation research on petrels and shearwaters’, Frontiers in Marine Science, 6, pp. 1–27. doi: 10.3389/fmars.2019.00094.

Rodríguez, A. et al. (2022) ‘Tracking Flights to Investigate Seabird Mortality Induced by Artificial Lights’, Frontiers in Ecology and Evolution, 9, p. 786557. doi: 10.3389/fevo.2021.786557.

Rodríguez, A. (2023) Conservation of Marine Birds. Elsevier. doi: 10.1016/C2020-0-03628-5.

Rodríguez, A., Dann, P. and Chiaradia, A. (2017) ‘Reducing light-induced mortality of seabirds: High pressure sodium lights decrease the fatal attraction of shearwaters’, Journal for Nature Conservation, 39, pp. 68–72. doi: 10.1016/j.jnc.2017.07.001.

Rodríguez, A., Rodríguez, B. and Negro, J. J. (2015) ‘GPS tracking for mapping seabird mortality induced by light pollution’, Scientific Reports, 5, pp. 1–11. doi: 10.1038/srep10670.

Rodríguez, B. et al. (2023) ‘Numbers of seabirds attracted to artificial lights should not be the only indicator of population trends’, Animal Conservation, pp. 2021–2023. doi: 10.1111/acv.12849.

Ruiz-Raya, F., Noguera, J. C. and Velando, A. (2022) ‘Light received by embryos promotes postnatal junior phenotypes in a seabird’, Behavioral Ecology, pp. 1–11. doi: 10.1093/beheco/arac079.

Ryan, P. G., Ryan, E. M. and Glass, J. P. (2021) ‘Dazzled by the light: the impact of light pollution from ships on seabirds at Tristan da Cunha’, Ostrich, 92(3), pp. 218–224. doi: 10.2989/00306525.2021.1984998.

Sánchez de Miguel, A. et al. (2021) ‘First estimation of global trends in nocturnal power emissions reveals acceleration of light pollution’, Remote Sensing, 13(16), pp. 1–12. doi: 10.3390/rs13163311.

Sánchez de Miguel, A. et al. (2022) ‘Environmental risks from artificial nighttime lighting widespread and increasing across Europe’, Science Advances, 8(37), pp. 1–10. doi: 10.1126/sciadv.abl6891.

Seltmann, M. W. et al. (2012) ‘Stress responsiveness, age and body condition interactively affect flight initiation distance in breeding female eiders’, Animal Behaviour, 84(4), pp. 889–896. doi: 10.1016/j.anbehav.2012.07.012.

Syposz, M. et al. (2018) ‘Factors influencing Manx Shearwater grounding on the west coast of Scotland’, Ibis, 160(4), pp. 846–854. doi: 10.1111/ibi.12594.

Syposz, M. et al. (2021) ‘Avoidance of different durations, colours and intensities of artificial light by adult seabirds’, Scientific Reports, 11(1), pp. 1–13. doi: 10.1038/s41598-021-97986-x.

Telfer, T. C. et al. (1987) ‘Attraction of hawaiian seabirds to lights: conservation efforts and effects of moon phase’, Wildlife Research, 15, pp. 406–413.

Troy, J. R. et al. (2013) ‘Using observed seabird fallout records to infer patterns of attraction to artificial light’, Endangered Species Research, 22(3), pp. 225–234. doi: 10.3354/esr00547.

Wilhelm, S. I. et al. (2021) ‘Effects of land-based light pollution on two species of burrow-nesting seabirds in Newfoundland and Labrador, Canada’, Avian Conservation and Ecology, 16(1), p. 12.

Witherington, B. and Bjorndal, K. A. (1991) ‘Influences of Wavelength and Intensity on Hatchling Sea Turtle Phototaxis: Implications for Sea-Finding Behavior’, Copeia, p. 1060. doi: 10.2307/1446101.

Zhao, X. et al. (2020) ‘Blue light attracts nocturnally migrating birds’, Condor, 122(2), pp. 1–12. doi: 10.1093/condor/duaa002.

